# The diversity of splicing modifiers acting on A_-1_ bulged 5’-splice sites reveals rules to guide rational design

**DOI:** 10.1101/2023.03.31.535100

**Authors:** Malard Florian, Antje Wolter, Julien Marquevielle, Estelle Morvan, Frédéric HT. Allain, Campagne Sebastien

## Abstract

Non-physiological alternative splicing patterns are associated with numerous human diseases. Among the strategies developed to treat these diseases, small molecule splicing modifiers are emerging as a new class of RNA therapeutics. The *SMN2* splicing modifier SMN-C5 was used as a prototype to understand their mode of action and discover the concept of 5’-splice site bulge repair. However, different small molecules harbouring a similar activity were also identified. In this study, we combined NMR spectroscopy and computational approaches to determine the binding modes of other *SMN2* and *HTT* splicing modifiers at the interface between U1 snRNP and an A_-1_ bulged 5’-splice site. Our results show that the other splicing modifiers interact with the intermolecular RNA helix epitope containing an unpaired adenine within a G_-2_A_-1_G_+1_U_+2_ motif, which is essential for their biological activity. We also determined structural models of risdiplam, SMN-CX, and branaplam bound to RNA, and solved the solution structure of the most divergent *SMN2* splicing modifier, SMN-CY, in complex with the RNA helix. These findings not only deepen our understanding of the chemical diversity of splicing modifiers that target A_-1_ bulged 5’-splice sites, but also identify common pharmacophores required for modulating 5’-splice site selection with small molecules.

## INTRODUCTION

Alternative pre-mRNA splicing is a crucial step in gene expression that determines mRNA localization, translation, and decay, and thus plays a significant role in regulating gene expression. Abnormal alternative splicing patterns have been associated with various human diseases (1).

A major limiting step in the splicing reaction is the definition of the 5’-splice site by the U1 small nuclear ribonucleoprotein (U1 snRNP), the first particle of the major spliceosome (2, 3). The ribonucleoprotein uses the 5’-end of the U1 snRNA to base pair with the 5’-splice site and initiates the splicing reaction (4). The recognition process is influenced by three factors: (i) the sequence of the 5’-splice site and its complementarity with the 5’-end of U1 snRNA (5, 6), (ii) the accessibility of the 5’-splice site, which may be trapped by inhibitory secondary structures (7), and (iii) the network of *trans*-acting splicing factors that can either enhance or inhibit U1 snRNP recruitment (8). Non-canonical base pair registers and bulged 5’-splice sites are often associated with alternative splicing patterns, and single nucleotide polymorphisms at the 5’-splice site have been linked to diseases (9–11). As a result, the definition of the 5’-splice site by U1 snRNP is a major target for splicing regulation and a key step for specific splicing correction. In recent decades, synthetic tools that enable the modulation of 5’-splice selection and the correction of alternative splicing have emerged and have begun to offer therapeutics for inherited diseases (12–16). There are currently two main classes of synthetic splicing switches: the antisense oligonucleotides (17) and the small molecule splicing modifiers (16, 18–20).

Antisense oligonucleotides modulate 5’-splice site selection indirectly, either by blocking key *cis*-regulatory RNA elements or disrupting the formation of inhibitory secondary structures (17), while small molecule splicing modifiers act directly at the interface between U1 snRNP and the 5’-splice site to modify 5’-splice site selection (19, 21). There are two classes of small molecule splicing modifiers that have been described: the *SMN2* and the *HTT* splicing modifiers, both of which act on A_-1_ bulged 5’-splice sites (16, 18). The mode of action of the *SMN2* splicing modifier family was recently proposed based on studies with the prototypical member, SMN-C5 (22). Upon treatment of human cells with SMN-C5, only a small number of exons responded, including *SMN2* exon 7. Analysis of their sequences showed an enrichment for a G_-2_A_-1_G_+1_U_+2_ motif at their exon-intron junctions. The solution structure of SMN-C5 bound to the RNA duplex formed by the U1 snRNA and the 5’-splice site (referred to as the RNA helix) revealed that the splicing modifier binds the RNA helix in the major groove at the exon-intron junction. Mechanistically, SMN-C5 acts as a molecular glue at the interface between U1 snRNP and the A_-1_ bulged 5’-splice site. The small molecule uses the carbonyl group of its central aromatic ring to form a direct hydrogen bond with the NH_2_ group of the unpaired adenine -1, explaining its specificity for 5’-splice sites containing a G_-2_A_-1_G_+1_U_+2_ motif. By stabilizing the unpaired adenine in the RNA base stack, SMN-C5 changes the geometry of the minor groove in the RNA helix, which stimulates and facilitates the binding of U1-C. SMN-C5 is not only a molecular glue between the 5’-splice site of *SMN2* exon 7 and the U1 snRNA, but also an allosteric effector of 5’-splice site definition on *SMN2* exon 7. This mechanism of action, which has not been seen before, has been referred to as 5’-splice site bulge repair, as the repair of the A_-1_ bulge leads to the strengthening of the 5’-splice site (Figure 1). While SMN-C5 specifically corrects *SMN2* exon 7 splicing using the bulge repair mechanism, it is unclear whether the molecule can represent the entire *SMN2* splicing modifier family (23–25), including the FDA-approved risdiplam, due to the chemical diversity of scaffolds among the family members.

**Figure 1.**
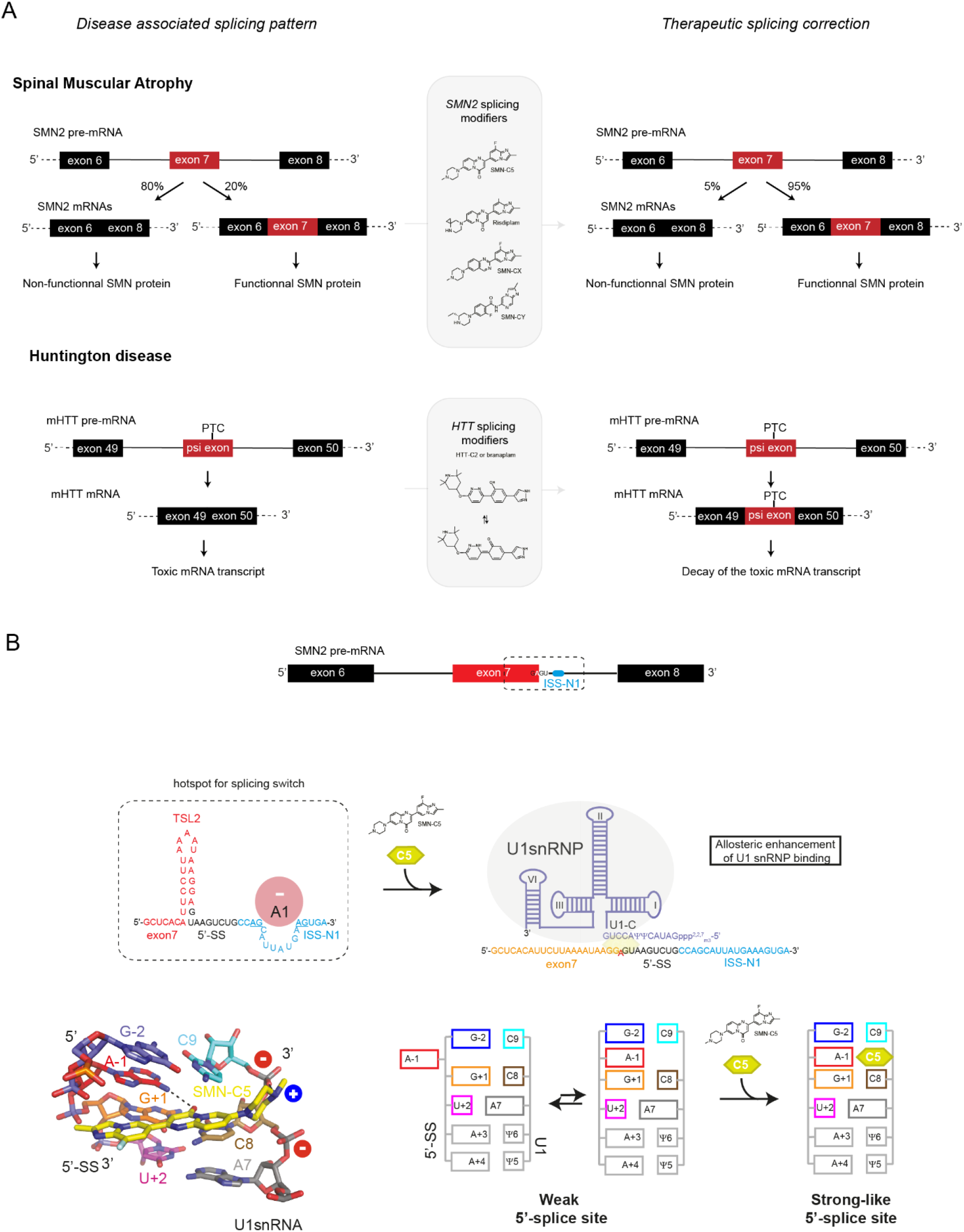
*SMN2* and *HTT* splicing modifiers: functions and mechanism of 5’-splice site bulge repair. A) Schematic representation of the splicing correction induced by *HTT* and *SMN2* splicing modifiers. Both classes of splicing modifiers promote specific exon inclusion. *SMN2* splicing modifiers induce *SMN2* exon 7 inclusion and allow the production of functional SMN protein from the *SMN2* gene. *HTT* splicing modifiers induce the inclusion of a pseudo exon (psi exon) containing a premature stop codon (PTC) triggering decay of the toxic *mHTT* mRNA. B) Mode of action of the *SMN2* splicing modifier SMN-C5. The hotspot for *SMN2* splicing correction is the weak 5’-splice site of exon 7 which is sequestered in an inhibitory secondary structure (TLS2), flanked by a strong splicing repressor (ISS-N1) and has a non-optimal sequence with an adenine in position -1. SMN-C5 stimulates the binding of U1 snRNP on this weak 5’-splice site by binding the RNA helix formed by U1 snRNP and the 5’-splice site at the exon-intron junction. SMN-C5 contacts specifically the unpaired adenine in position -1 and transforms the weak 5’-splice site of *SMN2* exon 7 into a stronger one.

*SMN2* splicing modifiers increase the inclusion of *SMN2* exon 7, thereby restoring the expression of the functional SMN protein from the *SMN2* gene and providing an orally available therapeutic for spinal muscular atrophy patients (23). In contrast, small molecule splicing modifiers that target the huntingtin mRNA (mHTT) reduce toxicity by promoting the inclusion of a PTC-containing poison exon that causes mRNA decay (18). These *HTT* splicing modifiers may represent potential therapeutics for Huntington’s disease as they lower the levels of toxic mHTT mRNA. Both classes of splicing modifiers identified so far promote exon inclusion, but have different functional consequences: *SMN2* splicing modifiers restore essential cellular function, while *HTT* splicing modifiers help cells get rid of toxic transcripts (Figure 1). Like *SMN2* splicing modifiers, HTT-lowering molecules also act on an A_-1_ bulged 5’-splice site (18) and are most effective when a purine-rich element is located in the exon.

Given that *HTT* splicing modifiers have different chemical scaffolds than SMN-C5, understanding how these molecules bind to the active site of the RNA may aid in optimizing lead compounds. Uncovering the structural diversity of splicing modifiers acting at the interface with U1 snRNA and an A_-1_ bulged 5’-splice site may also help to establish the rules governing their specificity for bulged splice sites, thereby enabling the rational design of small molecules that target other -1 bulged 5’-splice sites.

In order to understand how other *SMN2* and *HTT* splicing modifiers bind to the interface between U1 snRNP and the A_-1_ bulged 5’-splice site, we used a combination of *in vitro* binding assays, structure determination and computational approaches. Our atomic pictures of splicing modifier/RNA complexes showed that splicing modifiers representing three different chemotypes bind to the same epitope in the RNA helix, near an adenine in position -1 of the 5’ splice site required to achieve optimal activity. Through this work, we have gained valuable insights into the relationship between chemical scaffold diversity and the ability of splicing modifiers to target A_-1_ bulged 5’-splice in the context of disease.

## MATERIAL AND METHODS

### Sample preparation

The 5’-splice site of SMN2 exon 7 (5’-GGAGUAAGUCU-3’) and the U1 snRNA 5’-end (5’-AUACΨΨACCUG-3’) were purchased (Horizon), deprotected according to manufacturer instructions and lyophilized. The RNA helix was prepared by mixing in equimolar amounts each strand dissolved in the NMR buffer (MES d-4 5 mM pH 5.5, NaCl 50 mM), denatured at 75°C and annealed at room temperature. To reduce the amount of impurities, the sample was desalted using ZebaSpin column (Thermofisher). Risdiplam and branaplam were purchased (MCE Inhibitors) while SMN-CX and SMN-CY were provided by Roche. Risdiplam, branaplam and SMN-CX were resuspended in DMSO d8 while SMN-CY was soluble at high concentration in the NMR buffer. To collect NMR data on SMN-CY, a 1mM sample of SMN-CY was prepared in the NMR buffer. To collect the NMR data for the structure determination of the complex SMN-CY/RNA helix, the sample contained 1 mM of the RNA helix and 1.5 molar equivalent of SMN-CX in the NMR buffer. To collect NMR data on branaplam, a 10 mM sample was prepared in 100% DMSO d6 (Cambridge isotopes).

### NMR spectroscopy

The resonance assignment of SMN-CY was performed by combining 2D homonuclear experiments (TOCSY and NOESY) in 10% D_2_O. Data were recorded on a AVIII 750 MHz spectrometer (Bruker). The resonance assignment of branaplam was performed by combining 1D ^1^H, 1D ^13^C, 2D ^1^H-^1^H TOCSY, 2D ^1^H-^1^H NOESY, 2D ^1^H-^13^C HSQC and 2D ^1^H-^13^C HMQC. Data were collected on a AVIII 400 MHz and on a AVIII 700 MHz spectrometers (Bruker). In complex with SMN-CY, RNA resonance assignment was performed by combining 2D ^1^H-^1^H TOCSY, 2D ^1^H-^1^H NOESY and 2D ^13^C-^1^H natural abundance HSQC recorded on a AVIII 900 MHz spectrometer (Bruker). Previous assignment of the RNA helix served as a starting point. All the data were collected at 293K using Topspin (Bruker) and analysed with CARA (27).

### Structure calculations

To determine the structure of the RNA duplex-SMN-CY complex, we first determined the structure of the RNA within the complex. The RNA base pairing was determined based on the 2D ^1^H-^1^H NOESY recorded in 90% H_2_O. Angular restraints for the sugar pucker were derived from the analysis of the 2D ^1^H-^1^H TOCSY. Then, we combined this information with the chemical shifts and the experimental NOESY spectra recorded in 100% D2O (2D ^1^H-^1^H NOESY) for automatic NOE assignment and structure calculation using CYANA3.98.15 (27). A set of intramolecular NOE-derived distances was derived from the automatic analysis. In order to include SMN-CY in the CYANA calculation, a coordinate file together with a definition file for CYANA were generated using the Win!P software (http://www.biochem-caflisch.uzh.ch/download). Due to the excess of SMN-CY compared to the RNA duplex, strong NMR signals due to the presence of free SMN-CY impaired the automatic analysis of the intermolecular NOEs. Therefore, intermolecular NOE correlations were identified, integrated and calibrated manually. Initial structure calculations of the RNA duplex-SMN-CY complex were performed using CYANA in the torsion-angle space. The structures were further refined in the Cartesian space using the sander module of the AMBERTools22 package (28). The twenty lowest energy models extracted from the CYANA calculation were minimized, annealed and subjected to a short molecular dynamics step in water. The AMBER parameters for the SMN-CY were generated using the Antichamber module of AMBERTools22. Analysis of the calculation was performed using CYANA and structures were aligned and visualized using PyMol (Schrödinger).

### Docking procedures

We used the HADDOCK 2.4 algorithm to study the potential interactions between small molecules that modify splicing and the intermolecular RNA helix formed by the 5’ end of U1 snRNA and the A_-1_ bulged 5’-splice site (29). This algorithm can incorporate experimental data such as chemical shift perturbations or experimental distance restraints to predict the structure of a complex between RNA and drugs. We validated the binding pocket of these drugs within the RNA helix using NMR spectroscopy and found that the chemical shift perturbation profiles for all tested small molecule splicing modifiers were similar and consistent with the reference complex between SMN-C5 and the intermolecular RNA helix. Based on this information, we derived a set of distance restraints for use in docking calculations with HADDOCK using the solution structure of SMN-C5 in complex with the intermolecular RNA helix as a reference. These restraints included: (i) the primary amine group at A_-1_ forming a hydrogen bond with the small molecule, (ii) the positively charged amine in the piperazine group forming a salt bridge with a negatively charged oxygen in the phosphate group of C9 or C8, and (iii) the central planar unit of the drug inserting between the C8 and C9 C-H edges of the U1 snRNA. We used the bound conformation of the intermolecular RNA helix extracted from the solution structure of the duplex in complex with SMN-C5 (PDB code: 6HMO) as the target for docking simulations, and prepared the small molecules for docking using Marvin Sketch software to ensure their appropriate protonation states and 3D arrangement before exporting them in PDB format. The docking simulations were performed using the HADDOCK web server interface (30). The TBL files containing ambiguous and unambiguous distance restraints used in our study, as well as the corresponding ligand and RNA target PDB files, are described in the supplementary material.

## RESULTS

### Molecular basis for *SMN2* splicing correction by risdiplam

Risdiplam, a splicing modifier for spinal muscular atrophy, has undergone successful clinical evaluation and has been approved by regulatory agencies (23). It is the first oral treatment for this condition. Chemical modifications were made to the compound RG7800 (a homolog of SMN-C5) to improve its potency and reduce phototoxicity, metabolism, and basicity in the development of risdiplam (20). In comparison with SMN-C5, risdiplam differed in several ways: (i) the fluorine group on the first aromatic cycle was replaced with a methyl group; (ii) the methylated piperazine group was dealkylated to prevent N-dealkylation *in vivo*; and (iii) the piperazine group was modified by adding a cyclopropyl group at position 3 (Figure 2A). The modifications made to RG7800 in the creation of risdiplam had no effect on the mode of action compared to SMN-C5, risdiplam showed optimal biological activity when there was an adenine at position -1 of the 5’-splice site (31). Here, we used Nuclear Magnetic Resonance (NMR) spectroscopy to study the binding mode of risdiplam to the RNA helix formed by the 5’-splice site of SMN2 exon 7 and the 5’ end of U1 snRNA. When risdiplam was added to the RNA helix sample, we observed line broadening of the G+1 imino proton resonance and characteristic chemical shift perturbations of the H5-H6 TOCSY correlation peaks of C8 and U+2 (Figure 2B-C). These results suggest that risdiplam interacts with the RNA helix in a similar manner to SMN-C5. To get a more detailed understanding of how risdiplam binds to the RNA helix, we developed a docking protocol using a minimal number of restraints: (i) the primary amine group at A-1 must form a direct hydrogen bond with the small molecule, and (ii) the positively charged amine in the piperazine group must form a salt bridge with a negatively charged oxygen in the phosphate group of C9 or C8. The protocol was based on the SMN-C5/RNA helix complex and was validated using the experimentally determined structure (Supplementary Figure 1). The docking solutions were very similar to the NMR structure, and we used this approach to model risdiplam on the RNA helix. When performed with risdiplam, the docking calculation produced a cluster of low-energy solutions (Figure 2D). The lowest energy model of the RNA helix/risdiplam complex reveals that risdiplam interacts with the RNA helix in the major groove, in the pocket containing the GAGU motif (Figure 2E). As observed in the solution structure of SMN-C5 bound to the RNA helix, the central aromatic ring inserts between C8 and C9 of the U1 snRNA and forms a direct hydrogen bond with the amino group of the unpaired adenine at position -1. The positively charged amine in the piperazine group forms a salt bridge with the negatively charged oxygen in the phosphate group of C9 (Figure 2F). Like SMN-C5, risdiplam links the U1 snRNA and the 5’-splice site of SMN2 exon 7 and serves as the minimal *trans*-splicing factor (Figure 2G). Overall, our NMR spectroscopy and computational data suggest that risdiplam acts through the mechanism of 5’-splice site bulge repair.

**Figure 2.**
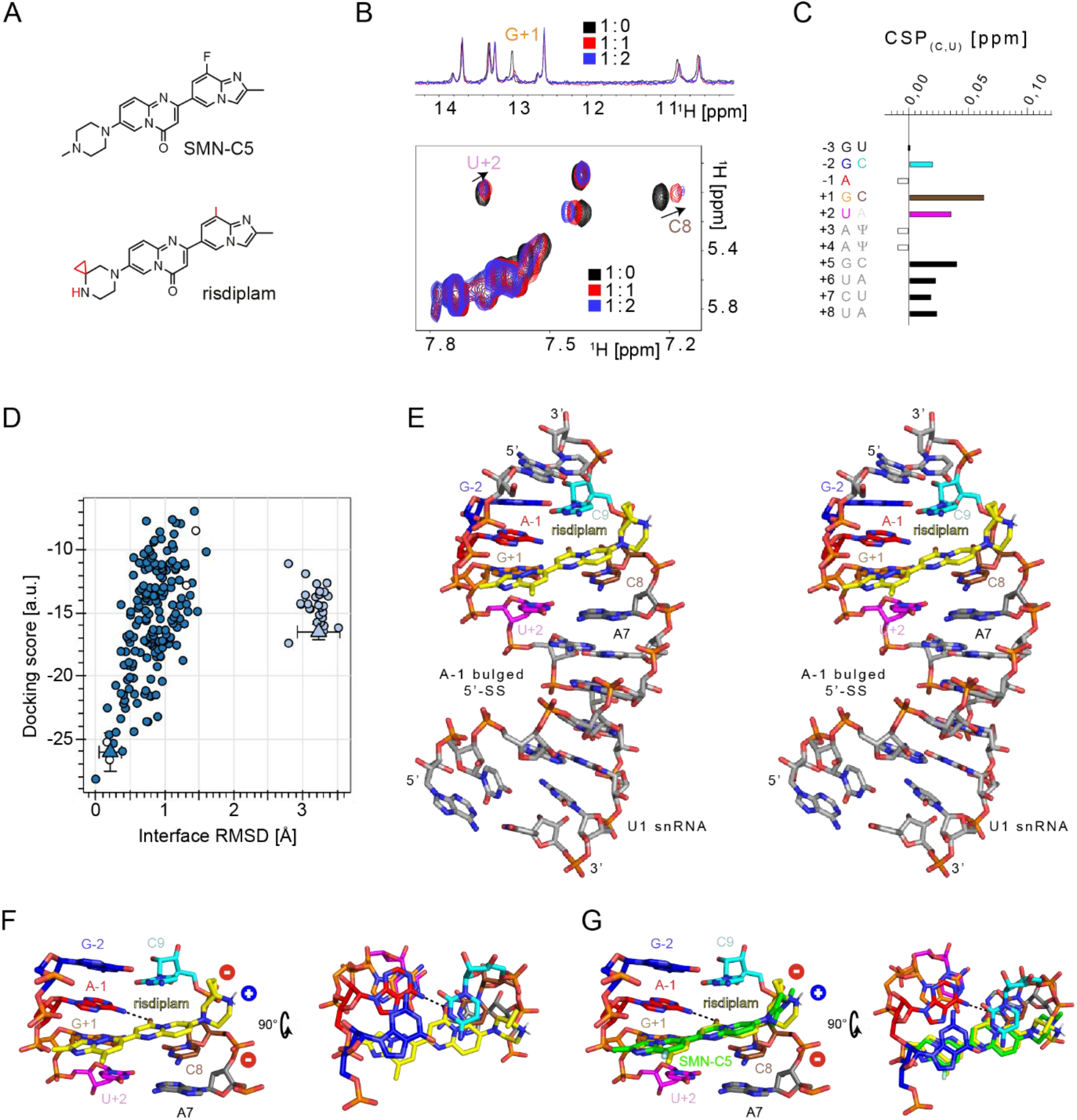
Molecular basis for *SMN2* splicing correction induced by risdiplam. A) Planar structures of SMN-C5 and risdiplam. The differences between both *SMN2* splicing modifiers are highlighted in red. B) Overlay of the 1D ^1^H (imino region) and 2D ^1^H-^1^H TOCSY (H6-H5 region) spectra of the RNA helix recorded upon successive addition of risdiplam. The spectra are coloured according the ratio RNA helix: risdiplam. C) Plot of the pyrimidine (H5-H6) chemical shift perturbations as a function of the positions on the 5’-splice site. White bars correspond to position without values (−1, +3 and +4). In position 7, the contributions of both pyrimidine has been added. D) Plot showing the correlation between the docking score (HADDOCK score) and the interface RMSD of the 200 docking solutions with respect to the lowest energy model. E) Stereoview of the lowest energy model of the complex formed by risdiplam and the RNA helix. F) Closed up views of the intermolecular interface between risdiplam and the RNA helix epitope. Intermolecular h-bonds are shown as dashed lines. G) Overlay of the lowest energy model and of SMN-C5 bound to the RNA helix (PDB ID 6HMO, structure 1). SMN-C5 is coloured in green.

### *SMN2* splicing correction can be achieved using a splicing modifier lacking the carbonyl group of the central aromatic ring

The structures of SMN-C5 and risdiplam bound to the RNA helix highlight the important role of the carbonyl group on the central aromatic unit (22). The carbonyl group provides specificity by forming a direct hydrogen bond with the primary amine of the unpaired adenine at position -1. Unlike SMN-C5 and risdiplam, the *SMN2* splicing modifier called SMN-CX lacks this carbonyl group (Figure 3A). Like SMN-C5 and risdiplam, and despite the absence of the carbonyl group on the central aromatic ring, SMN-CX showed optimal biological activity on the SMN2 system which has an adenosine at position -1 of the 5’-splice site (32). This finding was surprising and led us to verify whether SMN-CX binds to the RNA helix in the same way as SMN-C5 or risdiplam. Using NMR spectroscopy, we titrated the RNA helix with increasing amounts of SMN-CX and monitored the RNA resonances using 1D ^1^H and 2D ^1^H-^1^H TOCSY experiments (Figure 3B-C). Upon adding SMN-CX to the RNA helix sample, we observed the characteristic line broadening of the imino proton of G_+1_ and changes in the chemical shifts of the signals of C_8_ and U_+2_ on the TOCSY spectrum. These data suggest that SMN-CX binds to the RNA in the major groove near the exon-intron junction, similar to SMN-C5 and risdiplam. To get a more detailed view of the interaction between SMN-CX and the RNA helix, we used our docking protocol to generate structural models, assuming that the positive partial charge on the piperazine group would form a salt bridge with the phosphate group of C_8_ or C_9_ and that the adenine -1 amine group would be involved in a direct hydrogen bond with the small molecule. The docking approach was again very effective and produced a cluster of low-energy models (Figure 3D). The lowest energy model reveals that SMN-CX binds to the RNA helix in the major groove at the exon-intron junction (Figure 3E). The positive charge on the piperazine forms an electrostatic interaction with the phosphate group of C_9_, while the central aromatic ring inserts between C_8_ and C_9_ to position its azote atom to accept a hydrogen from the amino group of the unpaired adenine (Figure 3E). Overall, our experimental and computational data support a structural model for the interaction between SMN-CX and the RNA helix, demonstrating that the carbonyl group on the central aromatic unit of SMN-C5 and risdiplam is not necessary for specifically targeting A_-1_ bulged 5’-splice sites and can be replaced by other hydrogen bond acceptors.

**Figure 3.**
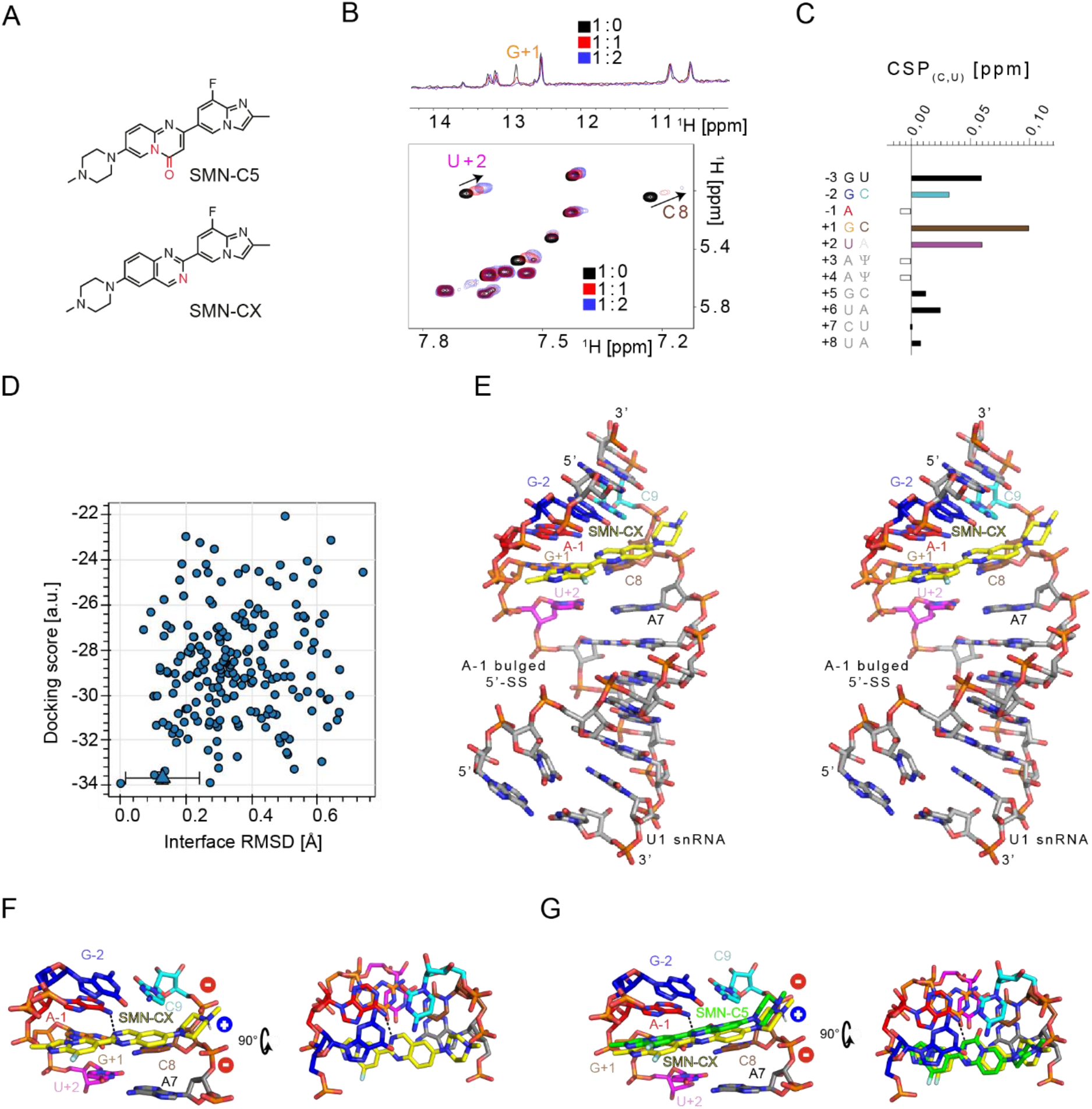
Molecular basis for SMN2 splicing correction induced by SMN-CX. A) Planar structures of SMN-C5 and SMN-CX. The differences between the two splicing modifiers are highlighted in red. B) Overlay of the 1D ^1^H (imino region) and 2D ^1^H-^1^H TOCSY (H6-H5 region) spectra of the RNA recorded upon successive addition of SMN-CX. The spectra are coloured according the ratio RNA helix : SMN-CX. C) Plot of the pyrimidine (H5-H6) chemical shift perturbations as a function of the positions on the 5’-splice site. White bars correspond to position without values (−1, +3 and +4). In position 7, the contributions of both pyrimidine has been added. D) Plot showing the correlation between the docking score (HADDOCK score) and the interface RMSD of the 200 docking solutions with respect to the lowest energy model. E) Stereoview of the lowest energy model of the complex formed by SMN-CX and the RNA helix. F) Closed up views of the intermolecular interface between SMN-CX and the RNA helix epitope. Intermolecular h-bonds are shown as dashed lines. G) Overlay of the lowest energy model and of SMN-C5 bound to the RNA helix (PDB ID 6HMO, structure 1). SMN-C5 is coloured in green.

### The size of the central aromatic unit can be reduced

The clinical evaluation of RG7800 (a homolog of SMN-C5) revealed a long-term safety issue with treatment. To address this problem, two approaches were taken: (i) optimization of RG7800, which led to the discovery and approval of risdiplam, and (ii) the search for other compounds with similar activity. This effort resulted in the identification, through rational drug design, of a new family of compounds with a benzamide core and a novel chemotype that can correct the splicing of *SMN2* exon 7 in a dose-dependent manner (24). During the structure-activity relationship study, a fluorine atom was added to the phenyl ring to create SMN-CY. The ^1^H resonances of SMN-CY were assigned using a combination of homonuclear NMR experiments (Supplementary Figure 2). The pyridyl analogue generates lone-pair repulsion between the pyridyl nitrogen and the carbonyl of the amide, as well as a favourable electrostatic interaction with the amide NH. In support of this, our NMR data showed that the amide proton of the molecule was strongly upfielded, indicating its involvement in a direct hydrogen bond with the fluorine atom (Figure 4A and Supplementary Figure 2). The presence of this weak interaction almost leads to a planar configuration of the carbonyl of the peptidyl-like bond and the benzamide. To understand how SMN-CY corrects *SMN2* splicing, we first determined whether SMN-CY interacts with the RNA helix (Figure 4B). Upon adding SMN-CY to the RNA helix sample, we observed characteristic chemical shift perturbations of the U_+2_ and C_8_ resonances and perturbations on the signal corresponding to the imino proton of G_+1_, suggesting that SMN-CY binds the same RNA pocket as SMN-C5 (Figure 4C). We also monitored changes on the RNA spectra using 2D ^1^H-^13^C HSQC experiments (Supplementary Figure 3). However, the perturbed RNA signals did not exhibit strong line broadening effects, as when the RNA helix was titrated with SMN-C5, risdiplam, or SMN-CX. This may be due to the reduction in size of the central aromatic ring and associated ring current effects. These data indicate that the size of the central aromatic unit can be reduced while still achieving *SMN2* splicing correction (24). To gain structural insights into the interaction between SMN-CY and the RNA helix, we attempted to solve the structure of the RNA/small molecule complex in solution.

**Figure 4.**
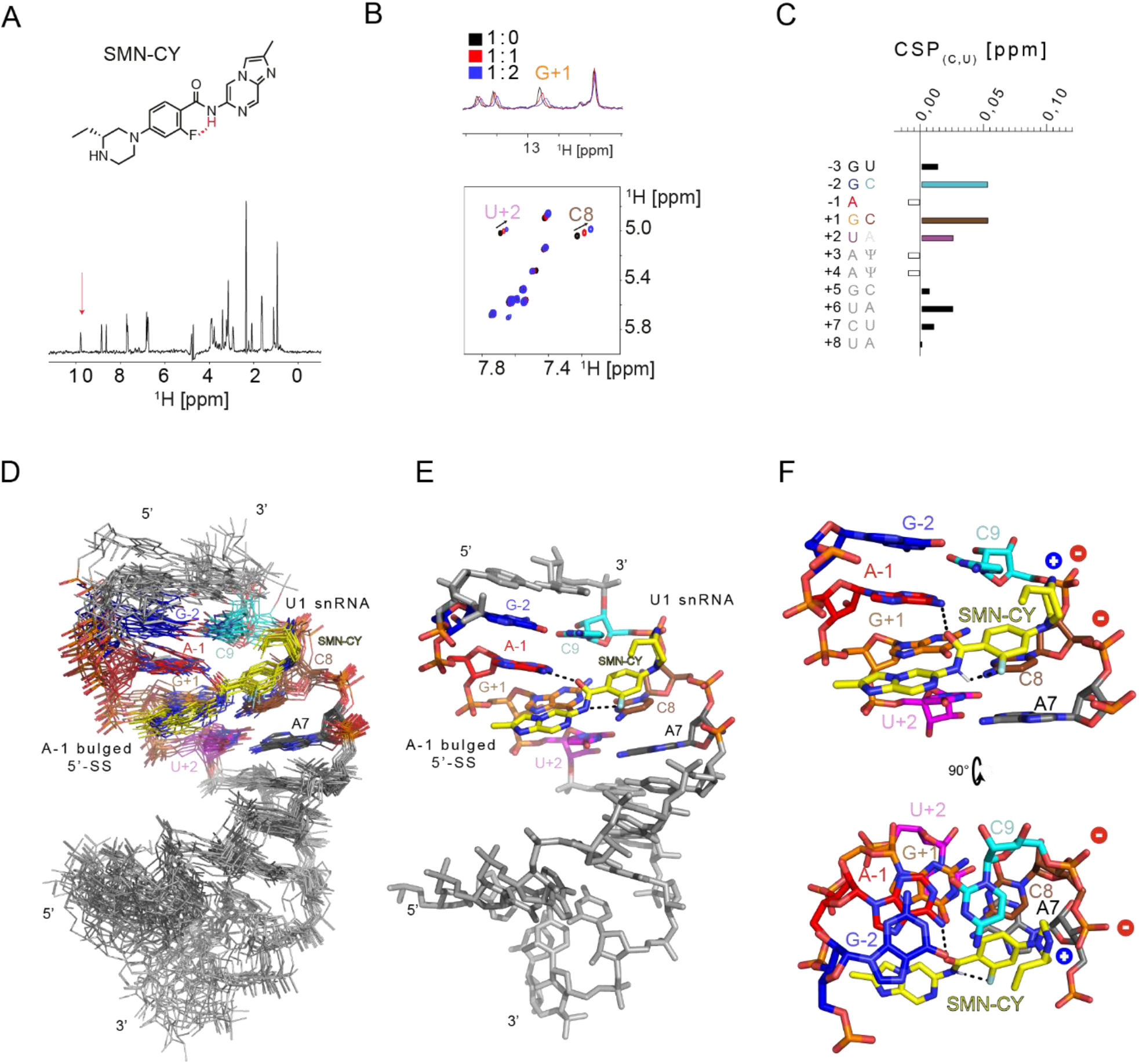
RNA binding activity of SMN-CY. A) Planar structures of SMN-CY and 1D 1H NMR spectra of SMN-CY recorded in 10% D_2_O. The signal that was assigned to the amide proton is shown by an arrow. B) Overlay of the 1D ^1^H (imino region) and 2D ^1^H-^1^H TOCSY (H6-H5 region) spectra of the RNA recorded upon successive addition of SMN-CY. The spectra are coloured according the ratio RNA helix : SMN-CY. C) Plot of the pyrimidine (H5-H6) chemical shift perturbations as a function of the positions on the 5’-splice site. White bars correspond to position without values (−1, +3 and +4). In position 7, the contributions of both pyrimidine has been added. D) Stereoview of the solution structure of SMN-CY bound to the RNA helix. E) Closed up view of the small molecule/RNA interface.

### Solution structure of SMN-CY bound to the RNA helix

To solve the structure of SMN-CY bound to the RNA helix, we assigned the proton resonances of SMN-CY (Supplementary Table 1) and utilized the previously assigned RNA resonances to solve the SMN-C5/RNA helix structure. We then prepared a sample containing the RNA helix in the presence of a 1.5 molar excess of SMN-CY and recorded 2D ^1^H-^1^H NOESY experiments with various mixing times. Using a mixing time of 200 ms, we identified intermolecular NOEs between SMN-CY and the RNA helix that defined the intermolecular interface (Supplementary Figure 4). Both methyl groups of the molecule provided clear intermolecular NOEs that oriented the small molecule within the major groove. On one side, Q28 was in close proximity to A_-1_ and U_+2_, while the ethyl branch on the piperazine was located closer to C_9_ and U_10_. The solution structure of the RNA helix bound to SMN-CY was determined using NOE-derived distances, including 18 intermolecular distances. The 14 NMR structures of the NMR ensemble superimpose well on the SMN-CY binding site with an average pairwise root mean square deviation of 0.88 ± 0.15 Å. The solution structure reveals that SMN-CY positions its piperazine positive partial charge near the phosphate group of C_9_ and inserts its benzamide between C_8_ and C_9_ aromatic rings. By interacting with the amide proton, the fluorine atom fixes the orientation of the carbonyl group of the peptidyl bond and places it in an optimal position to form a direct hydrogen bond with the amine group of the unpaired A_-1_. While SMN-CY represents a different chemotype than SMN-C5, their mode of action is very similar, and many common features were observed between both intermolecular interfaces, suggesting that these may be crucial for targeting the A_-1_ bulged 5’-splice site epitope.

### Molecular modelling of *HTT* splicing modifiers

Structural analysis of SMN-C/RNA complexes reveals that two different chemotypes can target a similar RNA epitope at the A_-1_ 5’-splice site. To increase our understanding of A_-1_ 5’-splice site targeting molecules, we decided to investigate a compound from a third chemotype, branaplam (also called HTT-C2). Originally developed as a SMN2 splicing modifier (33), clinical evaluation of branaplam was halted and it was recently repurposed as an efficient HTT splicing modifier (18, 34). This compound promotes the inclusion of a poison exon in the toxic mRNA mHTT and is being investigated as a potential therapeutic solution for Huntington disease. Like the SMN-C compounds, branaplam targets the A_-1_ bulged 5’-splice site and requires the presence of a purine-rich motif in the exon (18). Branaplam is based on a different chemical scaffold than the SMN-C compounds, consisting of a pyridazine substituted with a tetramethylated piperidin cycle on one side and a cyclohexa-2,4-dien-1-one attached to a piridinol. From a theoretical perspective, the small molecule exists in a tautomeric equilibrium between two forms (Figure 5A). In tautomer 1, a hydroxyl group is present on the cyclohexane moiety and the pyridazine cycle is non-protonated. In tautomer 2, a carbonyl group is present on the cyclohexane and the pyridazine cycle is protonated. The proton is shared between both aromatic cycles, making them coplanar and similar to the central ring of chemotype 1 (SMN-C5, risdiplam, and SMN-CX). To better understand the tautomeric equilibrium, we conducted NMR spectroscopy experiments in deuterated DMSO to observe exchangeable protons (Supplementary Figure 5). We assigned the resonances of the small molecules (Supplementary Table 2) and used a 2D ^1^H-^13^C HMQC to observe scalar coupling-mediated correlation between the protons of the cyclohexane ring and the carbon holding the hydroxyl group (tautomer 1) or the carbonyl group (tautomer 2). On the HMQC spectrum, we observed a correlation between the protons of the cyclohexane ring and a quaternary carbon at 158.4 ppm, corresponding to the carbon holding the hydroxyl group (Figure 5B). We did not observe a correlation between the protons of the cyclohexane and a potential ^13^C signal around 200 ppm that could correspond to the carbonyl group. In deuterated DMSO, we only observed tautomer 1, indicating that the equilibrium is skewed towards tautomer 1 in these conditions (Figure 5B). Unfortunately, similar experiments cannot be performed in aqueous solution or in the presence of RNA because we do not have access to isotopically labelled small molecules. Therefore, we investigated how branaplam/HTT-C2 interacts with the RNA helix. To monitor the binding of branaplam/HTT-C2 on the RNA helix, we recorded 1D ^1^H and 2D ^1^H-^1^H TOCSY spectra of the RNA helix upon the addition of branaplam. We observed typical chemical shift perturbations of C_8_, U_+2_, and the broadening of the G_+1_ imino protons, suggesting that branaplam/HTT-C2 targets the same RNA epitope as the SMN-C splicing modifiers (Figure 5C-D). It is also worth noting that the RNA helix contains another bulge downstream of the pseudouridine pair that is also partially occupied by branaplam, as indicated by additional chemical perturbations observed on the resonances of the pyrimidines from position +7, in line with our previous observations(21). As previously demonstrated in the context of HTT splicing correction, branaplam/HTT-C2 requires an adenine in position -1 to achieve optimal biological activity on the SMN2 system (18, 31). Overall, these data support the idea that branaplam/HTT-C2 binds to the same RNA epitope as other SMN-C compounds and acts through the mechanism of 5’-splice site bulge repair. To gain structural insights into the branaplam/RNA interaction, we used the docking protocol we previously established (Supplementary Figure 1). We defined ambiguous restraints to (i) position the partial positive charge held by the piperidine in a way that is compatible with the formation of a salt bridge with the phosphate of C_9_ or C_8_, and (ii) position the amine group of adenine -1 in a direct hydrogen bond with branaplam. We also enforced proton sharing between the oxygen atom of the cyclohexane and the nitrogen of the pyridazine. The docking calculation converged well and revealed a solution that explains how this third chemotype can target the A_-1_ bulged 5’-splice site (Figure 5E). The structural model of branaplam bound to the RNA helix showed that the tetramethylpiperidine interacts with the phosphate group of C_9_ through a salt bridge and that the central aromatic ring is inserted between C_8_ and C_9_. By sharing its exchangeable proton with the adjacent nitrogen of the pyridazine, the resulting large flat entity positions its oxygen atom to form a direct hydrogen bond with the amine group of the unpaired adenine. This explains how branaplam/HTT-C2 can correct the splicing of target exons using the 5’-splice site bulge repair mechanism. In conclusion, branaplam/HTT-C2 is a third chemotype that can correct the splicing of A_-1_ bulged 5’-splice sites by targeting the same RNA epitope as chemotypes 1 and 2.

**Figure 5.**
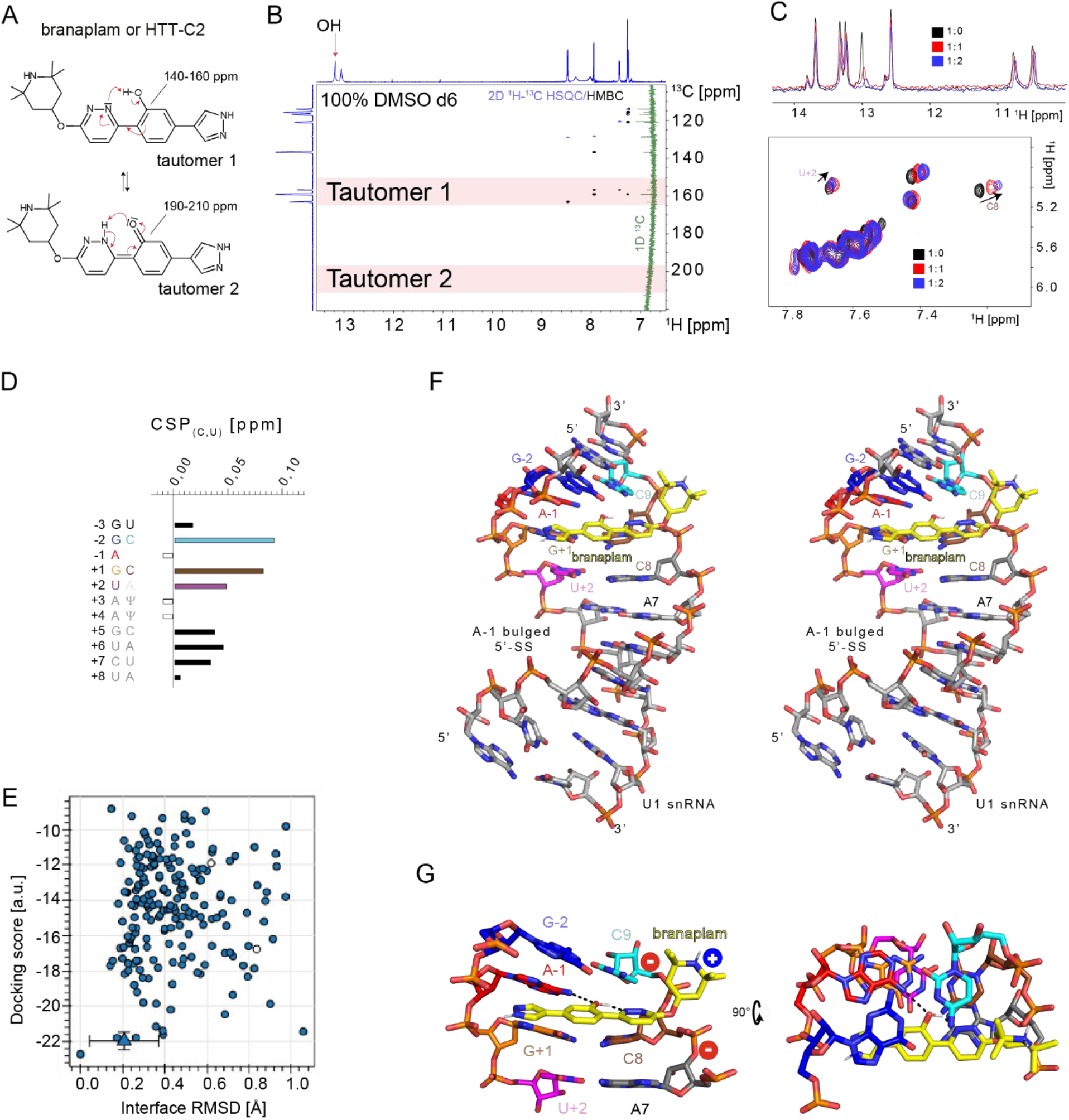
Structural basis for branaplam/HTT-C2 splicing correction. A) Planar structures of branaplam/HTT-C2. The small molecule is predicted to exchange between two tautomeric forms. B) Overlay of the 1D ^13^C (green), 2D ^1^H-^13^C HSQC (blue) and 2D ^1^H-^13^C HSQC (black) of branaplam/HTT-C2 recorded in 100% d6-DMSO. C) Overlay of the 1D ^1^H (imino region) and 2D ^1^H-^1^H TOCSY (H6-H5 region) spectra of the RNA recorded upon successive addition of branaplam. The spectra are coloured according the ratio RNA helix : branaplam. D) Plot of the pyrimidine (H5-H6) chemical shift perturbations as a function of the positions on the 5’-splice site. White bars correspond to position without values (−1, +3 and +4). In position 7, the contributions of both pyrimidine has been added. E) Plot showing the correlation between the docking score (HADDOCK score) and the interface RMSD of the 200 docking solutions with respect to the lowest energy model. E) Stereoview of the lowest energy model of the complex formed by branaplam/HTT-C2 and the RNA helix. F) Closed up view of the branaplam/RNA interface.

**Figure 6.**
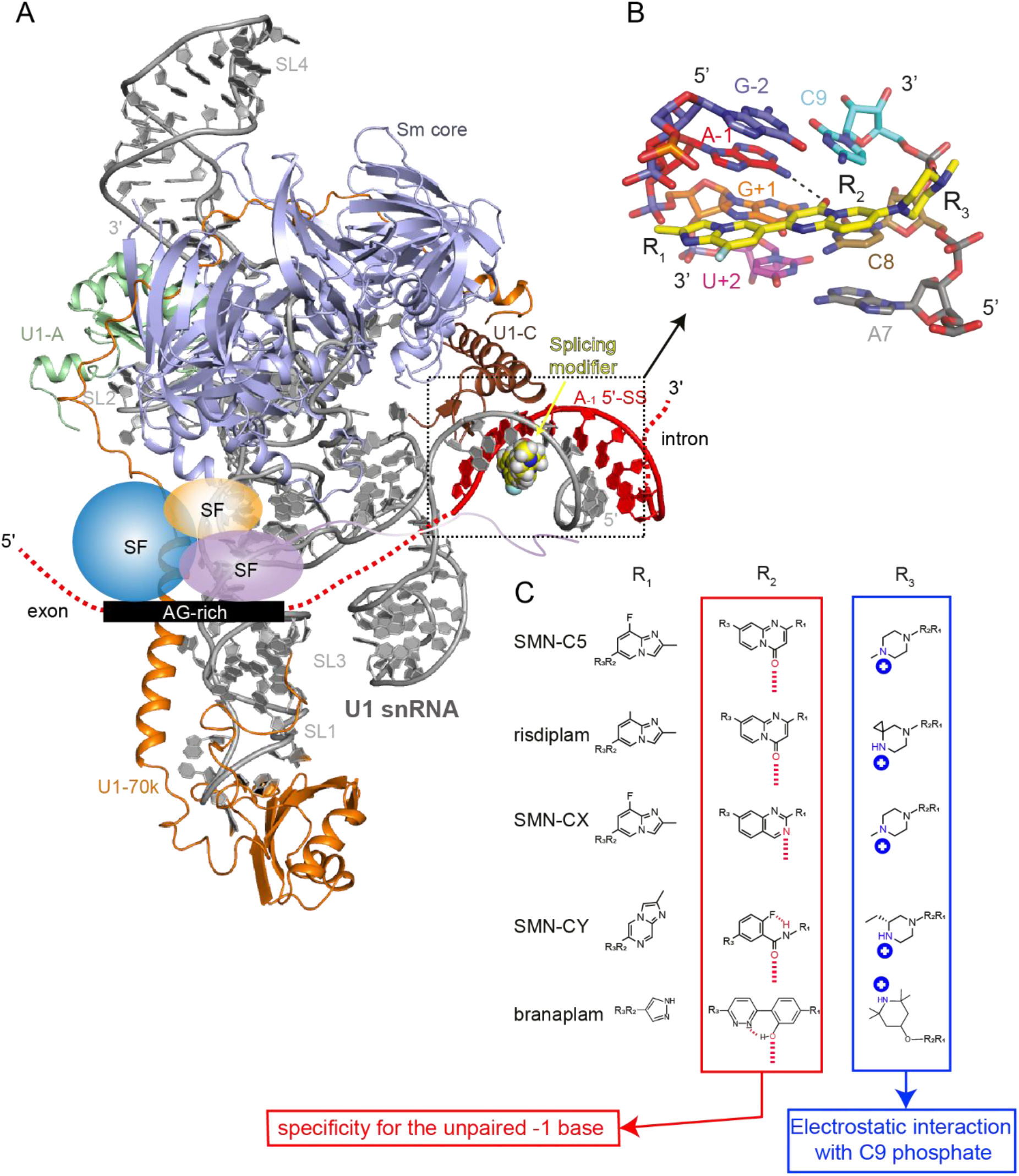
Chemical diversity of splicing modifiers acting on A-1 bulged 5’-splice site. A) Model of U1 snRNP bound to an A-1 5’-splice site in presence of SMN-C5. The exon position was extrapolated in order to depict the AG-rich motif and the splicing factors that are recruited. B) Closed up view of the RNA helix epitope bound by SMN-C5. The splicing modifier has been split in three functional parts: R_1_, R_2_ and R_3_. C) Functional analysis of the composition and role of splicing modifiers acting on A-1 bulged 5’-splice site.

## DISCUSSION

Single nucleotide polymorphisms at 5’-splice sites have been linked to numerous human diseases (9, 11) and are documented in the DBASS databases (35). Diseases are often caused by mutations at the invariant G_+1_ nucleotide and at positions -1, +2, and +5. In this study, we examined the chemical diversity of splicing modifiers targeting A_-1_ bulged 5’-splice sites for specific splicing correction, a promising class of RNA therapeutics (2). While the SMN2 splicing modifier SMN-C5 served as the prototype for this family and helped to clarify the mechanism of action and the concept of “5’-splice site bulge repair” (22), several small molecule splicing modifiers with different chemical scaffolds have similar biological activities (20, 24, 33). We selected four molecules representing three chemotypes and their ability to bind to the RNA:RNA interface between U1 snRNP and an A-1 bulged 5’-splice site within a G_-2_A_-1_G_+1_U_+2_ motif. Our data show that the four additional molecules bind to the same RNA epitope close to the unpaired adenosine at position -1. By providing atomic views of the small molecules’ RNA interfaces, we identified several common features. The small molecules cross the entire major groove and consist of three units: one in contact with the U1 snRNA (A), the central aromatic unit (B), and one in contact with the 5’-splice site (C). Unit A is often a piperazine or piperidine moiety that is positively charged at physiological pH and forms a salt bridge with the phosphate backbone of the U1 snRNA. Removing the charge or substituting the terminal amine greatly reduces biological activity (21). The basicity of the SMN-C series was modulated by modifying the substitution of the piperazine, which was crucial for the development of risdiplam (20). Piperidine or piperazine moieties provide flexible scaffolds that are suitable for this task, but do not provide specificity. Unit B is the central component of the molecule and provides specificity for the unpaired adenine. It inserts between the C8 and C9 C-H edges, a hydrophobic patch on the RNA epitope. This is why unit B must be aromatic and planar. More importantly, unit B often contains a carbonyl group that directly contacts the amine group of A-1. We have shown that the carbonyl is not essential for specificity and can be replaced by another hydrogen bond acceptor, as we observed with SMN-CX. Our structural models generally show that the carbonyl group must be in a fixed position to contact adenine -1. It is either embedded in an aromatic ring (SMN-C5, risdiplam) or stabilized by weak interactions (SMN-CY and branaplam). For SMN-CY and branaplam/HTT-C2, the rigidification of the carbonyl group position is particularly unique. In the case of SMN-CY, the carbonyl group is embedded in an amide bond attached to a fluoro-benzene. As previously proposed during SMN-CY optimization, the amide bond is fixed in the same plane as the aromatic ring by the formation of a direct hydrogen bond between the fluorine and the amide proton, leading to a strongly upfielded chemical shift of the amide proton. This weak interaction is essential for placing the carbonyl group in an ideal orientation to establish a direct hydrogen bond with the unpaired adenine and ensure optimal biological activity. The example of branaplam/HTT-C2 is also interesting because the small molecule is involved in a tautomeric exchange in which the hydroxyl group held by the cyclohexa-2,4-dienol forms a direct hydrogen bond with a nitrogen atom of the pyridazine. As a result, both aromatic rings are coplanar and this weak interaction locks the oxygen atom in an ideal position to interact with the unpaired adenine. The central ring controls the specificity for the unpaired base. Finally, unit C is often an aromatic ring with nitrogen atoms, but its role is unclear. In our structures, unit C is close to the AGU motif and may provide additional stacking, polar interactions or water mediated contacts that are extremely difficult to observe using solution state NMR spectroscopy. It could be interesting to add arms to this cycle to contact the phosphate backbone.

Overall, the diversity of splicing modifiers that act on A_-1_ bulged 5’-splice sites reveals rules for guiding rational design and expanding the repertoire of small molecule splicing modifiers. While our structural characterization of splicing modifiers bound to the RNA helix provides valuable insights for rational design of new small molecules, it does not explain why among the 200,000 A_-1_ bulged 5’-splice sites in the genome (33), SMN2 or HTT splicing modifiers are so selective and alter the splicing of only a few exons. Notably, both types of splicing modifiers require an A_-1_ bulged 5’-splice site as well as a purine-rich element in the exon (21, 33). While the role of the A_-1_ bulged 5’-splice site has been explained through the mechanism of 5’-splice site bulge repair, the role of the purine-rich element remains unclear. We previously showed that the purine-rich element acts as an enhancer of the splicing correction (22) and proposed that *trans*-splicing factors bound by the purine-rich element could help stabilize U1 snRNP on weak 5’-splice sites. However, it cannot be ruled out that splicing factors bound to the purine-rich element directly stabilize the RNA helix formed upon 5’-splice site recognition and also contact the small molecule to provide additional specificity. Future rational design efforts to alter the base specificity at position -1 of small molecule splicing modifiers will need to take this criterion into account.

## AVAILABILITY

Not applicable.

## ACCESSION NUMBERS

NMR chemical shifts and atomic coordinates for the reported solution structure have been deposited with the BioMagResBank (BMRB) under the accession code 34784 and the Protein Data bank under accession number 8CF2.

## SUPPLEMENTARY DATA

Supplementary Data are available at NAR online.

## ACKNOWLEDGEMENTS

We would like to thank the NMR platforms of ETH Zurich and of the European Institute of Chemistry and Biology (IECB). We would like to thank Roche Basel for providing the compounds SMN-CX and SMN-CY.

## TABLE AND FIGURES LEGENDS

**Table 1.**
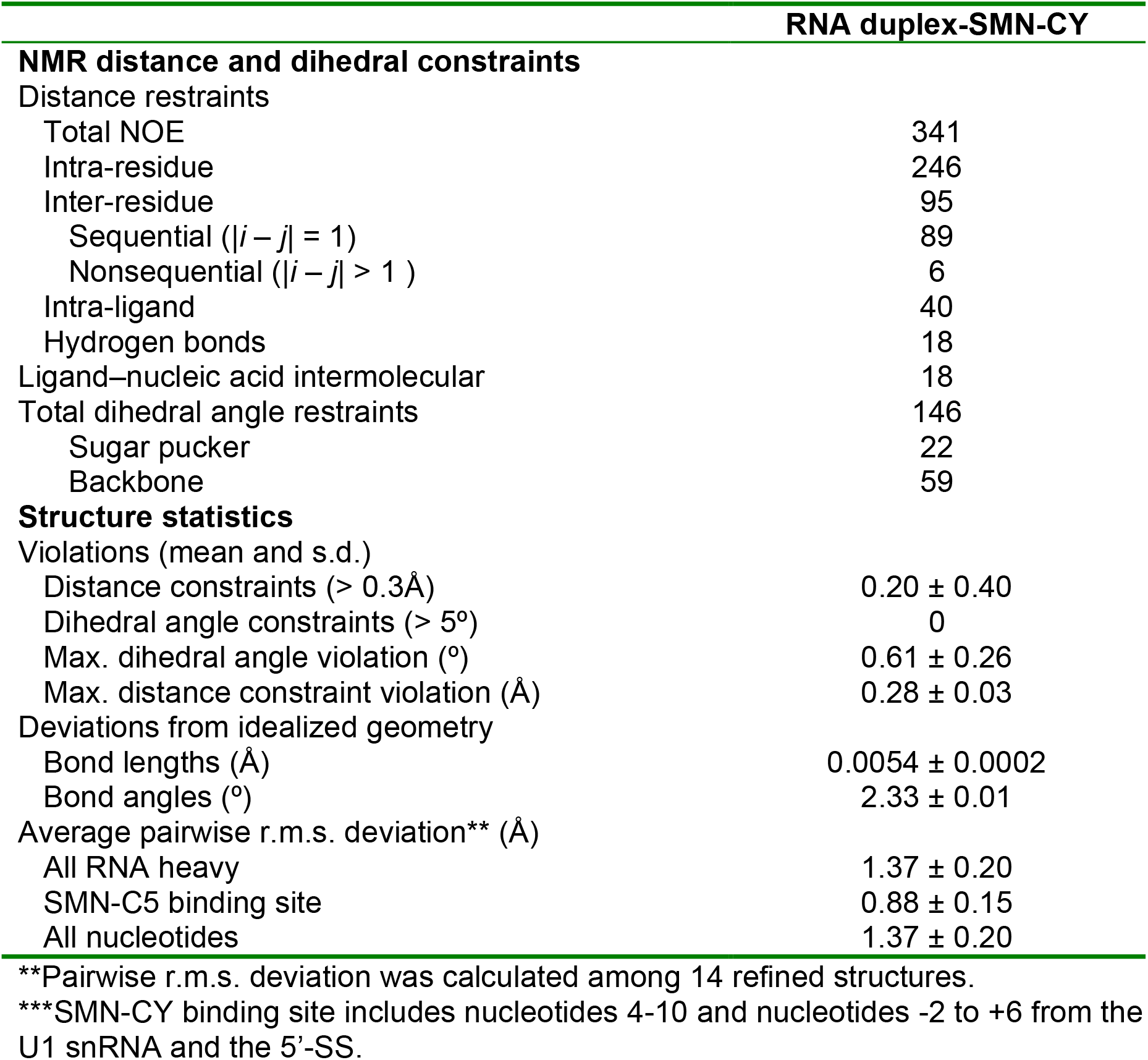
NMR and refinement statistics for the RNA duplex in complex with SMN-CY

