## supplementary file for "The diversity of splicing modifiers acting on A_-1_ bulged 5’-splice sites reveals rules to guide rational design"

Supplementary data checklist:

- Supplementary Figure 1. Development of the docking procedure on the interaction between SMN-C5 and the RNA helix.
- Supplementary Figure 2. Resonance assignment of SMN-CY.
- Supplementary Figure 3. Monitoring the interaction between SMN-CY and the RNA duplex.
- Supplementary Figure 4. Identification of intermolecular NOE between SMN-CY and the RNA duplex.
- Supplementary Figure 5. Resonance assignment of branaplam.
- Supplementary Table 1. SMN-CY chemical shifts at 293K.
- Supplementary Table 2. Branaplam chemical shifts at 293K in 100% DMSO-d6.

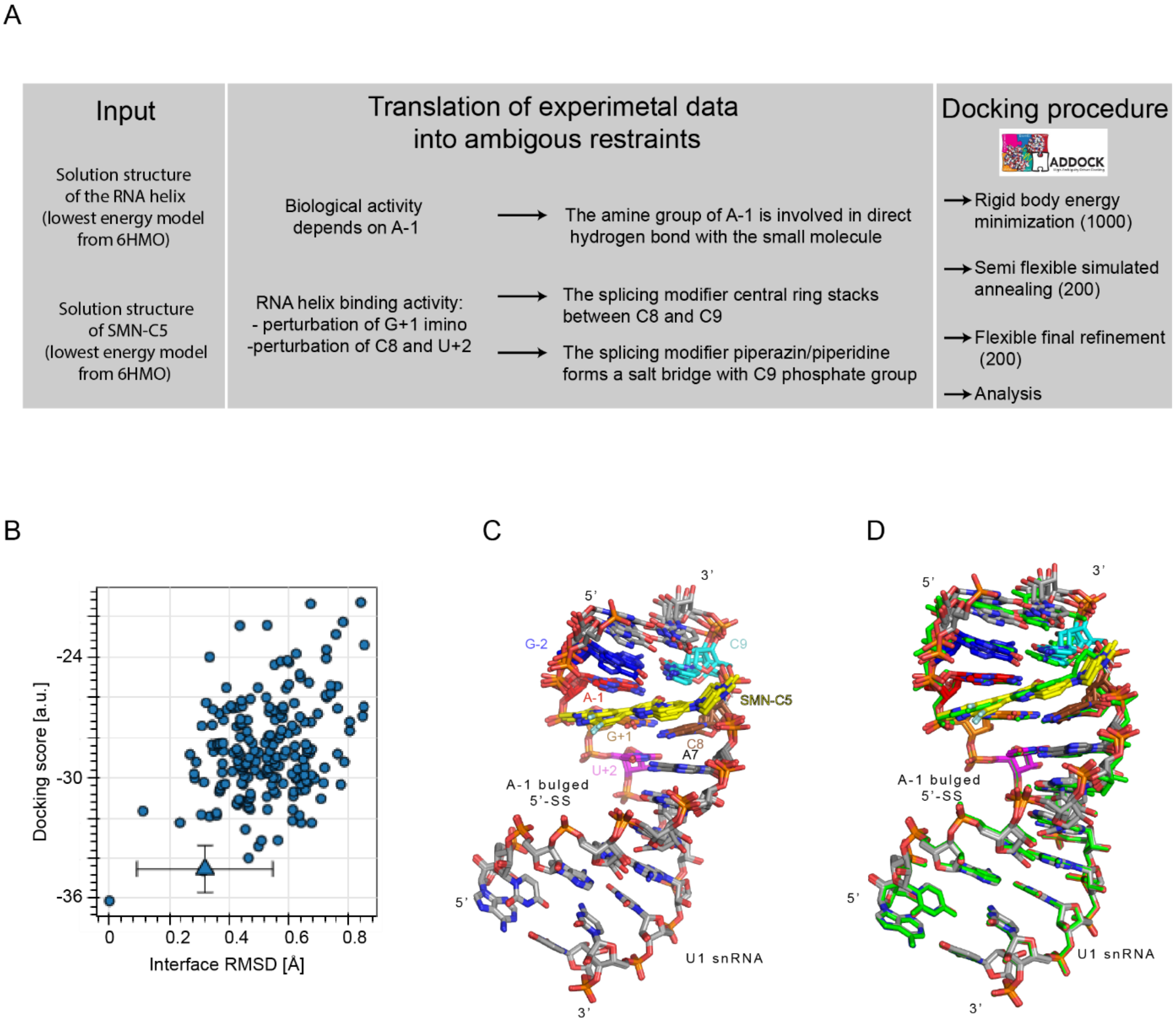


**Supplementary Figure 1. Development of the docking procedure on the interaction between SMN-C5 and the RNA helix.** A) Description of the NMR data driven docking approach using the HADDOCK 2.4 web server. B) Plot showing the correlation between the docking score (HADDOCK score) and the interface RMSD of the 200 docking solutions with respect to the lowest energy model of the complex RNA duplex / SMN-C5. C) Overlay of the four lowest energy models of the docking calculation. D) Superimposition of the four lowest energy models of the docking calculation and the lowest energy model of the solution structure of the RNA duplex bound by SMN-C5 (PDB ID 6HMO).

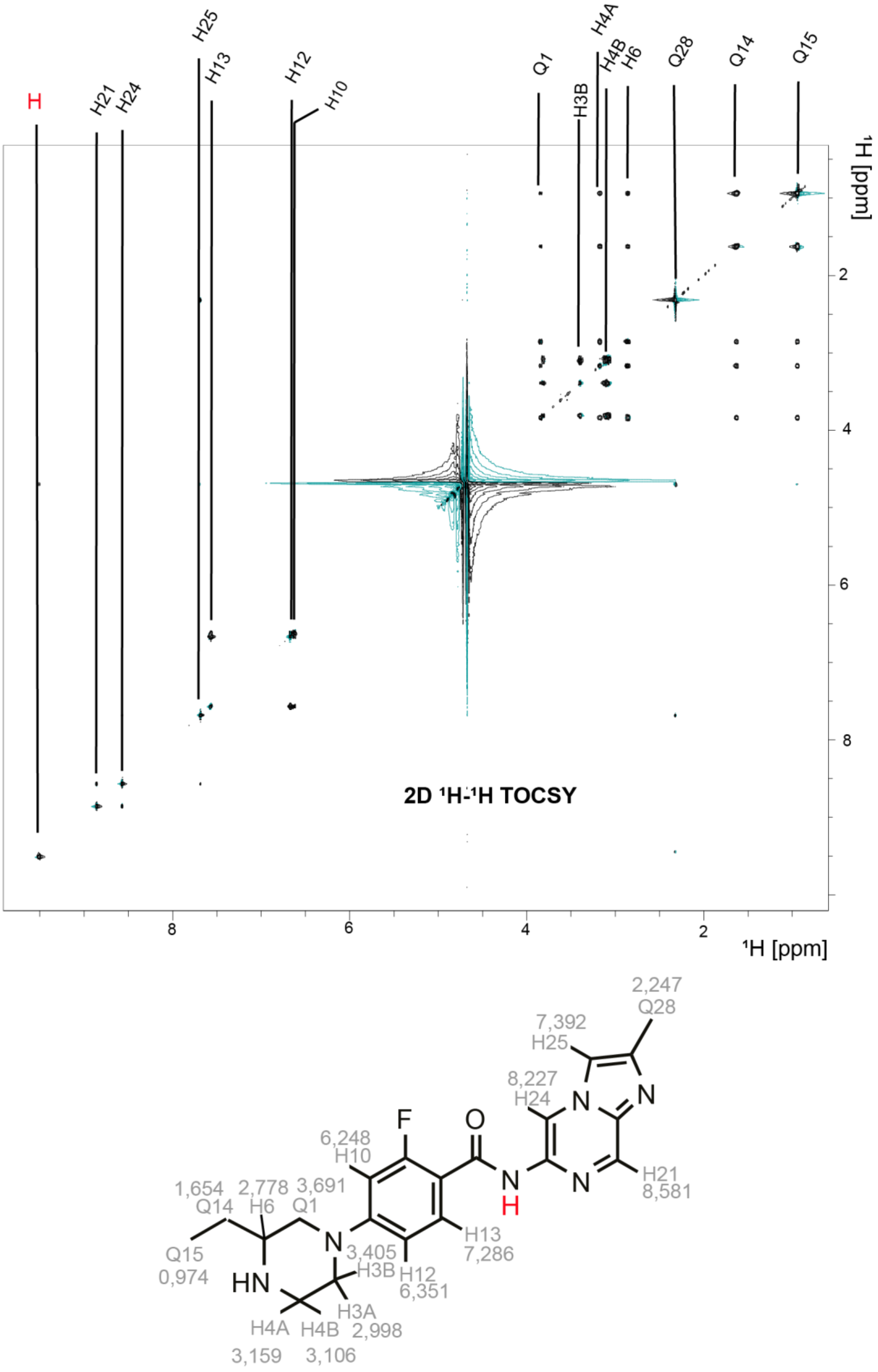


**Supplementary Figure 2. Resonance assignment of SMN-CY.** A) 2D $^{1}H$-$^{1}H$ TOCSY spectrum of SMN-CY. Below, the chemical structure of SMN-CY is shown.

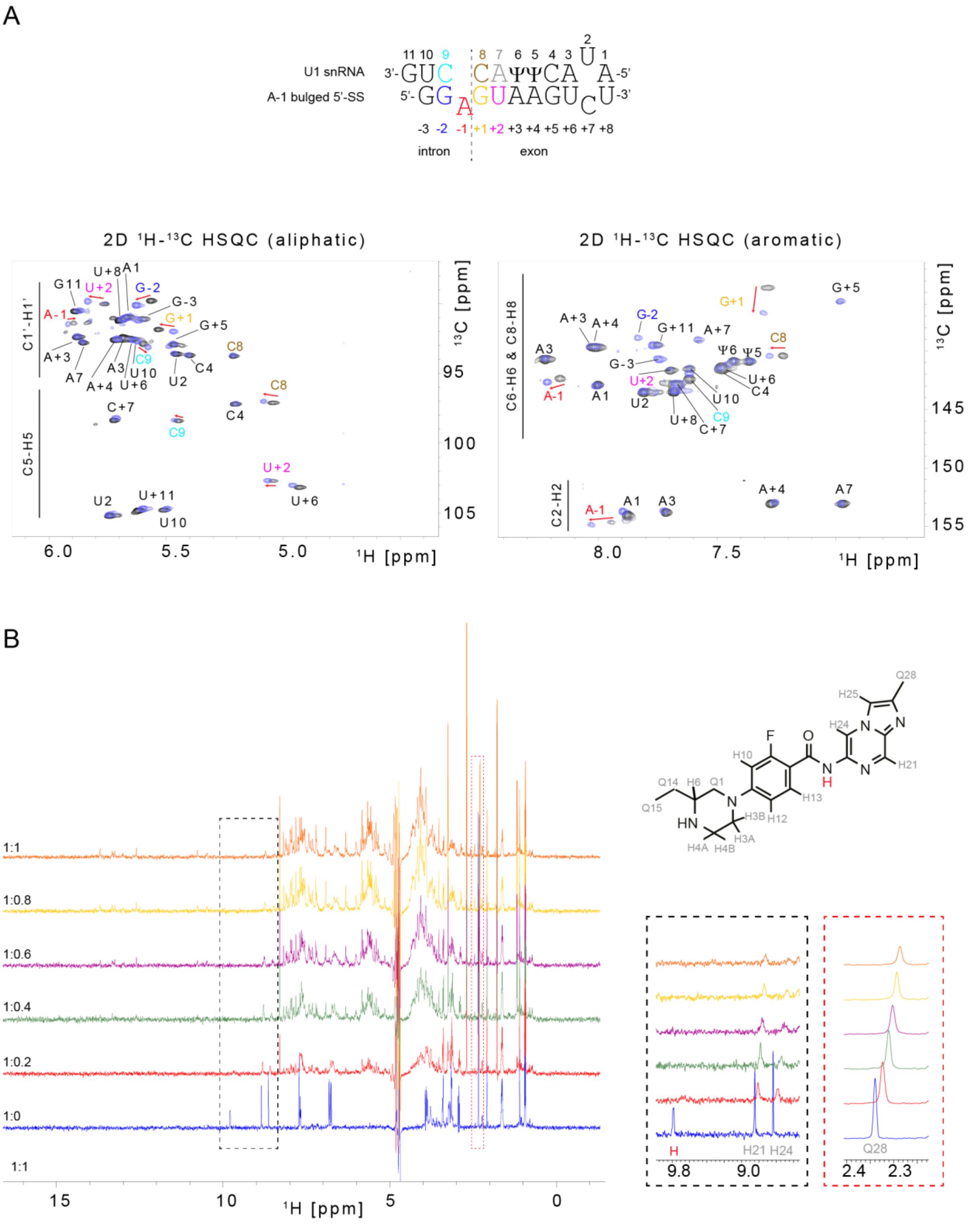

**Supplementary Figure 3. Monitoring the interaction between SMN-CY and the RNA duplex.** A) Schematic representation of the sequence of the RNA duplex. Below, the 2D $^{1}H$-$^{13}C$ HSQC spectra of the RNA duplex before (black) and after addition (blue) of SMN-CY. B) titration of SMN-CY by the RNA duplex followed by 1D $^{1}H$ NMR spectroscopy.

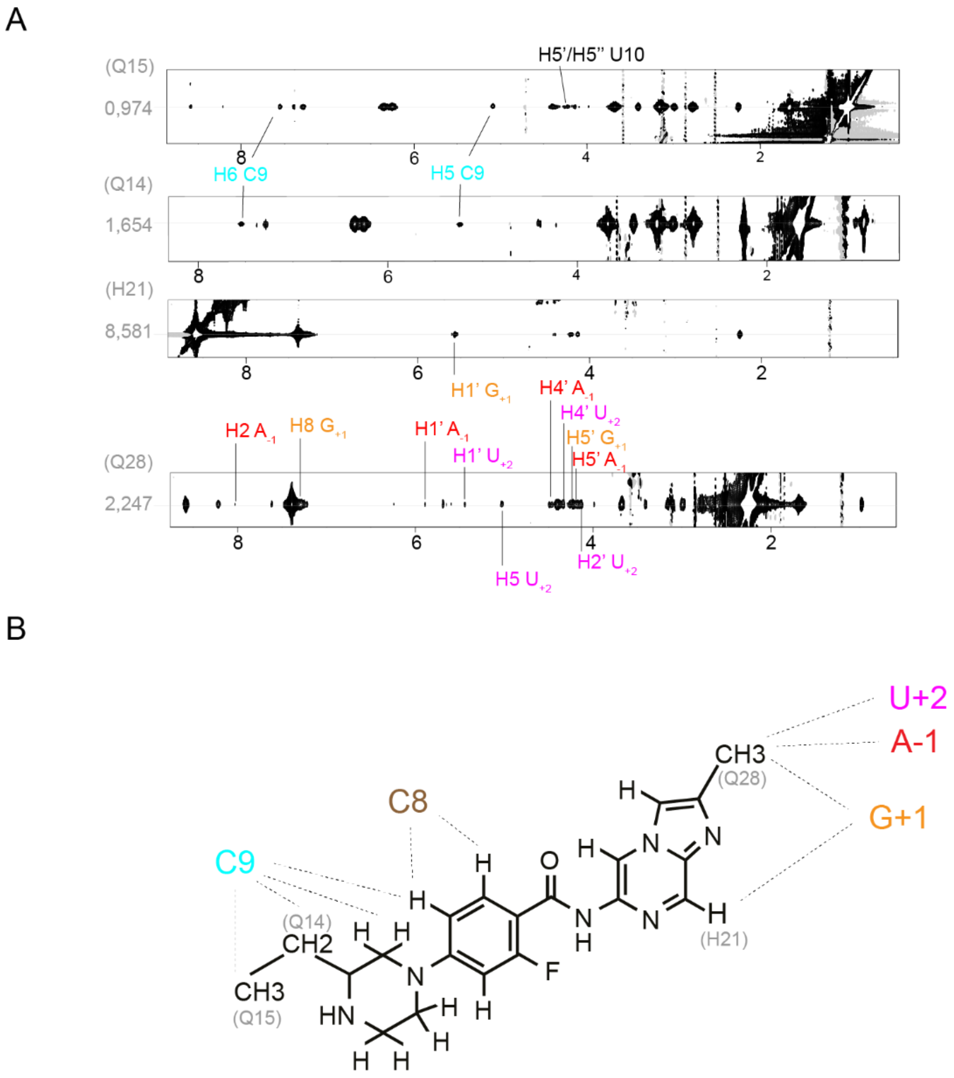


**Supplementary Figure 4. Identification of intermolecular NOE between SMN-CY and the RNA duplex.** A) Portion of the 2D $^1$H-$^1$H NOESY spectrum showing intermolecular NOEs between SMN-CY and the RNA duplex. B) Schematic representation of the proximity between the bases and SMN-CY.

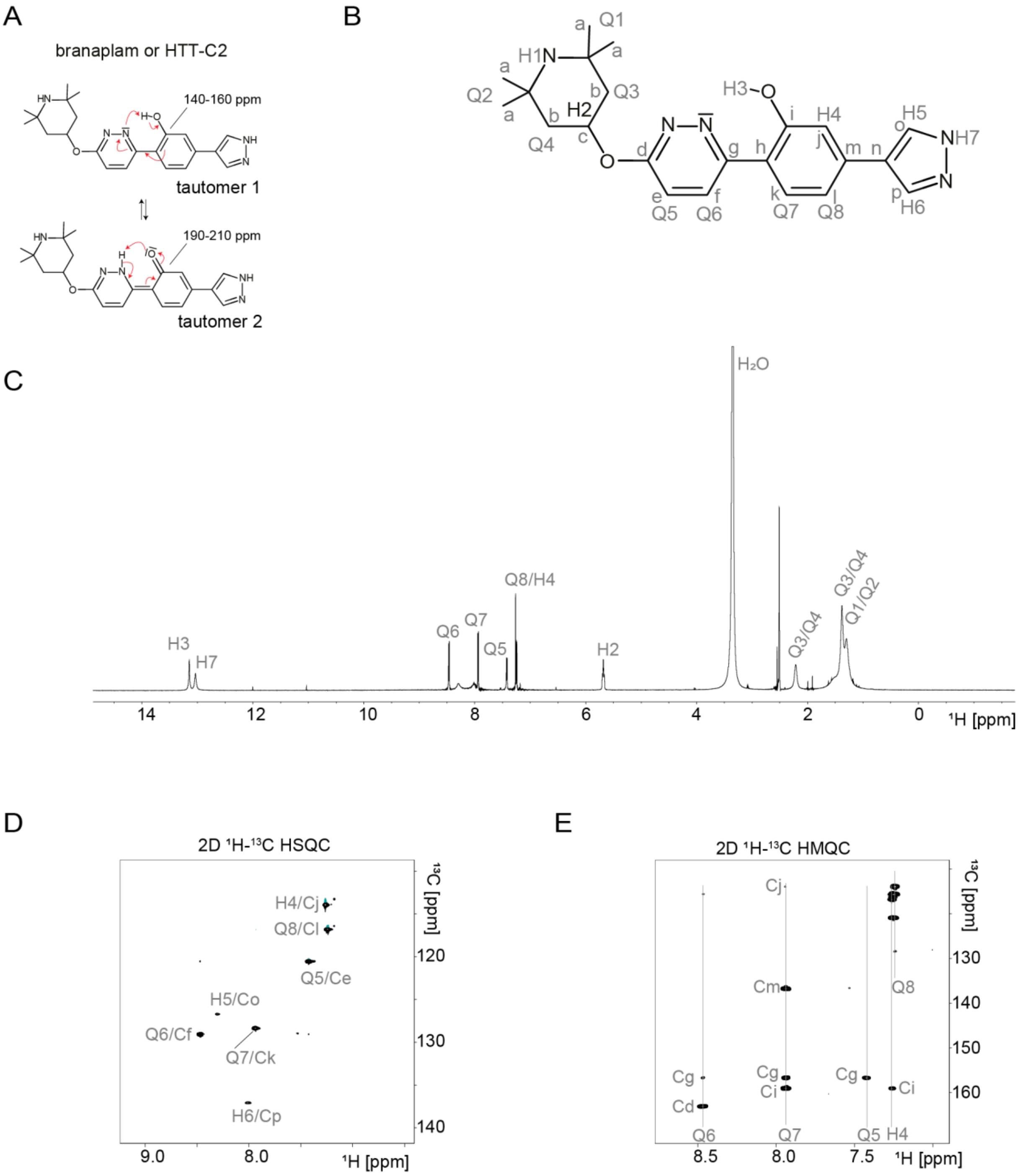


**Supplementary Figure 5. Resonance assignment of branaplam.** A) Schematic representation of the tautomeric equilibrium of branaplam. The planar structures of the two tautomeric forms are shown. B) Planar structure of the alcohol form of branaplam with atom names. C) 1D $^{1}H$ NMR spectrum of branaplam recorded in 100% DMSO-d6. D) Portion of the 2D $^{1}H$-$^{13}C$ HSQC spectrum of branaplam recorded in 100% DMSO-d6. E) Portion of the 2D $^{1}H$-$^{13}C$ selective HMQC spectrum of branaplam recorded in 100% DMSO-d6.

**Supplementary Table 1. SMN-CY chemical shifts at 293K.**

| Atom name* | Chemical shift free (ppm) | Chemical shift RNA-bound (ppm) |
|---|---|---|
| Q14 | 1.625 | 1.654 |
| Q15 | 0.935 | 0.974 |
| Q28 | 2.247 | 2.339 |
| H6 | 2.864 | 2.778 |
| Q1 | 3.819 | 3.681 |
| H3A | 2.873 | 2.988 |
| H3B | 3.397 | 3.405 |
| H4A | 3.175 | 3.159 |
| H4B | 3.090 | 3.106 |
| H10 | 6.761 | 6.248 |
| H12 | 6.818 | 6.351 |
| H13 | 7.680 | 7.286 |
| H21 | 8.849 | 8.581 |
| H24 | 8.638 | 8.227 |
| H25 | 7.716 | 7.392 |
| H (amide) | 9.780 | Broad, undetermined |

*Atom names are defined in Supplementary Figure 3B.

**Supplementary Table 2. Branaplam chemical shifts at 293K in 100% DMSO-d6.**

| Atom name* | Chemical shift free (ppm) |
|---|---|
| Q1** | 1.29/1.38 |
| Q2** | 1.29/1.38 |
| Q3** | 1.51/2.22 |
| Q4** | 1.51/2.22 |
| Q5 | 7.43 |
| Q6 | 8.48 |
| Q7 | 7.95 |
| Q8 | 7.25 |
| H2 | 5.68 |
| H3 | 13.18 |
| H4 | 7.27 |
| H5 | 8.30 |
| H6 | 8.03 |
| H7 | 13.05 |
| Ca | 27.51 |
| Cb | 41.53 |
| Cc | 69.94 |
| Cd | 162.63 |
| Ce | 120.09 |
| Cf | 128.63 |
| Cg | 156.26 |
| Ch | 120.46 |
| Ci | 158.62 |
| Cj | 113.49 |
| Ck | 127.92 |
| Cl | 116.37 |
| Cm | 136.28 |
| Cn | 115.18 |
| Co | 126.38 |
| Cp | 136.28 |

*Atom names are defined in Supplementary Figure 5.

**Stereospecific assignment was not performed.
